# The effect of global signal regression on DCM estimates of noise and effective connectivity from resting state fMRI

**DOI:** 10.1101/634063

**Authors:** Hannes Almgren, Frederik Van de Steen, Adeel Razi, Karl Friston, Daniele Marinazzo

## Abstract

The influence of the global BOLD signal on resting state functional connectivity in fMRI data remains a topic of debate, with little consensus. In this study, we assessed the effects of global signal regression (GSR) on effective connectivity within and between resting-state networks – as estimated with dynamic causal modelling (DCM) for resting state fMRI (rsfMRI). DCM incorporates a forward (generative) model that quantifies the contribution of different types of noise (including global measurement noise), effective connectivity, and (neuro)vascular processes to functional connectivity measurements. DCM analyses were applied to two different designs; namely, longitudinal and cross-sectional designs. In the modelling of longitudinal designs, we included four extensive longitudinal resting state fMRI datasets with a total number of 20 subjects. In the analysis of cross-sectional designs, we used rsfMRI data from 361 subjects from the Human Connectome Project. We hypothesized that (1) GSR would have no discernible impact on effective connectivity estimated with DCM, and (2) GSR would be reflected in the parameters representing global measurement noise. Additionally, we performed comparative analyses of the informative value of data with and without GSR. Our results showed negligible to small effects of GSR on connectivity within small (separately estimated) RSNs. For between-network connectivity, we found two important effects: the effect of GSR on between-network connectivity (averaged over all connections) was negligible to small, while the effect of GSR on individual connections was non-negligible. Contrary to our expectations, we found either no effect (in the longitudinal designs) or a non-specific (cross-sectional design) effect of GSR on parameters representing (global) measurement noise. Data without GSR were found to be more informative than data with GSR; however, in small resting state networks the precision of posterior estimates was greater using data after GSR. In conclusion, GSR is a minor concern in DCM studies; however, individual between-network connections (as opposed to average between-network connectivity) and noise parameters should be interpreted quantitatively with some caution. The Kullback-Leibler divergence of the posterior from the prior, together with the precision of posterior estimates, might offer a useful measure to assess the appropriateness of GSR, when nuancing data features in resting state fMRI.

## 1. Introduction

The fMRI signal is corrupted by noise from several sources; for example, motion-induced noise, background (thermal) noise, and non-neural physiological noise that arises from cardiac and respiratory processes (Liu, 2016). This is especially a problem in resting state fMRI research, which usually aims to quantify low-frequency fluctuations in the absence of explicit perturbations. Extensive research has been carried out to develop and apply methods to de-noise the (resting-state) fMRI signal (e.g., Erdoğan et al., 2016; Kasper et al., 2017; Rummel et al., 2013). Some studies focus on modeling explicit (external) sources of noise (e.g., by including motion and cardiac signal as regressors in a GLM; see, e.g., Kasper et al., 2017), others focus on identifying signal and noise directly from fMRI signals (e.g., using ICA; see, e.g., Rummel et al., 2013). A widely applied (group of) method(s) – used to denoise fMRI signals – is to correct resting-state fMRI time-series for fluctuations in the global signal (GS), which is the average signal across all voxels of the entire MRI volume. Different types of corrections for GS fluctuations have been developed and applied, including GS regression (GSR), GS normalization, and GS subtraction (Liu, Nalci, & Falahpour, 2017).

The use (or omission) of GSR in fMRI connectivity studies has been a hot topic of debate (Murphy & Fox, 2016). The method has commonly been applied to account for multiple non-neural sources of noise (e.g., motion and cardiac-induced signal). Some studies have shown that GSR might increase the efficiency of detecting significant functional connectivity (Liu et al., 2017). Fox et al. (2009), for example, showed that global signal regression (GSR) enhanced both the spatial specificity of positive correlations and detection of anti-correlation between networks (e.g., between default mode and dorsal attention networks). Similarly, Yan et al. (2013) showed that GSR decreased the correlation between motion and functional connectivity. On the other hand, other studies have shown that global signal regression introduces spurious connectivity, and leads to complex region-dependent biases (Anderson et al., 2011; Murphy et al., 2009; Saad et al., 2012). Murphy et al. (2009), for example, showed both analytically and through simulations that global signal regression causes spurious negative connectivity. Saad et al. (2012) and Gotts et al. (2013) showed that group differences can become distorted after GSR. With this background, some authors have argued for new perspectives on how to study GSR (see, e.g., Power et al., 2017; Uddin, 2017).

Typically, studies investigating the impact of global signal corrections have focused on measures of connectivity that do not include a biologically plausible (forward) model; for example, correlation and independent component analysis (ICA). Therefore, connectivity estimates in these studies are not separated from estimates of hemodynamic processes or measurement noise. Dynamic causal modelling (DCM; Friston, Harrison, & Penny, 2003) is a method that allows such separation, by explicitly incorporating parameters representing state and measurement noise, effective connectivity and (neuro)vascular processes. Parameters representing noise are separated into three components: neural fluctuations that drive the system (i.e., state noise), observation or measurement noise (both local and global; e.g., caused by changes in scanner temperature), and sampling error (caused by imperfect sampling). Parameters representing (neuro)vascular processes include parameters modelling vasodilatory signal decay (related to neurovascular coupling), mean blood transit time (the average time it takes for blood to pass the veins), and the ratio of intravascular to extravascular contributions to the measured fMRI signal (Stephan et al., 2007).

In the present study, we assessed the effects of global signal regression on effective connectivity and noise parameters as estimated by (spectral) DCM for resting state fMRI (Friston, Kahan, Biswal, & Razi, 2014; Razi, Kahan, Rees, & Friston, 2015). We expected that (1) global signal regression would not have a substantial impact on effective connectivity, and (2) its main effect would be reflected in the parameters representing global observation noise. In addition, we investigated whether data with GSR affords a greater information gain compared to data without GSR – as well as increasing the precision of posterior connectivity estimates. To address these questions, we decomposed free energy a lower bound on the log model evidence – into an accuracy and a complexity term. The latter reflects the amount of information gain afforded by the data.

Four resting state networks were analysed: three resting state networks (RSNs; namely somatomotor, saliency, and default mode network) and an additional larger network comprising all three networks. To allow for sufficient generalizability, and ensure a comprehensive examination of how GSR affects connectivity, we performed analyses using two different designs; namely, longitudinal and cross-sectional designs. For the former, we used four longitudinal datasets (see Almgren et al., 2018), to which a hierarchical approach was applied (i.e., from sessions to subjects, and from subjects to group). The benefits of such a hierarchical analysis are that: (a) the total variance is captured by multiple variance components (e.g., between-session variance, between-subject variance), hence rendering parameter estimates potentially more precise, (b) within-subject effects (e.g., fluctuations in the amount of noise) are mitigated, and (c) conclusions are not limited to one acquisition protocol or dataset. The second design was cross-sectional, for which we used the human connectome project’s dataset. The benefits of this dataset are that (a) data are preprocessed by a standardized HCP pipeline, (b) the sample size is large (361 unrelated subjects), hence mitigating subject-specific effects.

## 2. Methods

### 2.1 Datasets and subjects

The longitudinal datasets were acquired by four different research institutions (see, Choe et al., 2015, Filevich et al., 2017, Gordon et al., 2017, and Laumann et al., 2015). Together they comprised 20 subjects (11 females, mean and standard deviation of age at onset study: 30.1 ± 5.2) and contained a total of 653 rsfMRI sessions (at the least 10 for each subject). For a further description of the longitudinal datasets, see Almgren et al. (2018).

In addition, the Human Connectome Project’s 900 subject release (HCP; Van Essen et al., 2012) was analysed as cross-sectional dataset. To avoid issues arising with data from related subjects, we only modelled data from 361 unrelated subjects (194 females; mean and standard deviation age: 28.7 ± 3.7). Only the first session of all subjects was analysed.

### 2.2 Data Analyses

All analyses were performed using the SPM12 software package (revision 6906; Wellcome Centre for Human Neuroimaging; www.fil.ion.ucl.ac.uk/spm/software/spm12), including DCM for resting state fMRI (DCM12; revision 6801), and parametric empirical Bayes (PEB; revision 6778).

#### 2.2.1 Preprocessing

Concerning the longitudinal datasets, the same preprocessing and time series extraction steps were used as in Almgren et al. (2018). In short, the initial five images for each resting state fMRI session were discarded, then rsfMRI data were corrected for differences in slice time (using the central slice as a reference), realigned to the first volume of each session, coregistered to an anatomical image (anatomical image prior to first functional scan session), normalized to MNI space and smoothed using a Gaussian kernel (FWHM = 6mm). Three sessions (across all subjects and datasets) were discarded because of insufficient quality. Concerning the cross sectional data from HCP dataset, the minimally preprocessed rsfMRI data of the first scanning session of each subject were used (see, Glasser et al., 2013). These data were additionally smoothed using a Gaussian kernel of 6mm FWHM.

#### 2.2.2 Time-series extraction

Extraction of regional time-series was the same for both types of datasets – and is described in detail in Almgren et al. (2018). In short, voxels showing low frequency fluctuations were identified using a GLM with a discrete cosine basis set as regressor of interest (0.0078–0.1Hz; where the number of components was a function of the number of scans and TR) and eight nuisance regressors (six head motion regressors, CSF signal from 5mm ROI in circulatory system, WM signal from 7mm ROI in brainstem). For the analyses with GSR the average time-series across the whole brain (using spm_global.m) were included as a nuisance regressor (see paragraph 2.2.5). The SPM produced by specifying and estimating an F-contrast across all DCT components was masked with ROIs from template ICA maps (Smith et al., 2009). Table 1 shows the exact coordinates for each region; Figure 1 shows the regions superimposed on a template brain. Time-series were computed as the principal eigenvariate of signals centred on the peak voxel of the SPM (sphere radius = 8mm) in each ROI. This procedure allowed for subject-specific peak locations of low-frequency fluctuations within the boundaries of the (subject independent) template ROIs. The somatomotor network (SMR), salience network (SAL), and default mode network (DMN), comprised respectively three, five, and four regions. The same time-series were also included in the combined network (excluding mPFC because of its proximity to ACC).

**Table 1.** ROI regions and coordinates. Coordinates were adopted from template ICA images (Smith et al., 2009). *Not included in combined network.

| Network | Region | Coordinates |  |  |
| --- | --- | --- | --- | --- |
| Somatomotor | Supplementary motor area | 0 | -12 | 50 |
|  | Right precentral gyrus | 44 | -16 | 48 |
|  | Left postcentral gyrus | -38 | -26 | 56 |
| **Salience** | Anterior cingulate cortex | 0 | 3 | 22 |
|  | Left middle frontal gyrus | -28 | 54 | 14 |
|  | Right middle frontal gyrus | 30 | 52 | 14 |
|  | Left insula | -36 | 12 | -2 |
|  | Right insula | 32 | 20 | 4 |
| **Default mode** | Precuneus | 2 | -58 | 30 |
|  | Medial prefrontal cortex* | 2 | 56 | -4 |
|  | Left inferior parietal cortex | -44 | -60 | 24 |
|  | Right inferior parietal cortex | 54 | -62 | 28 |

**Figure 1.**
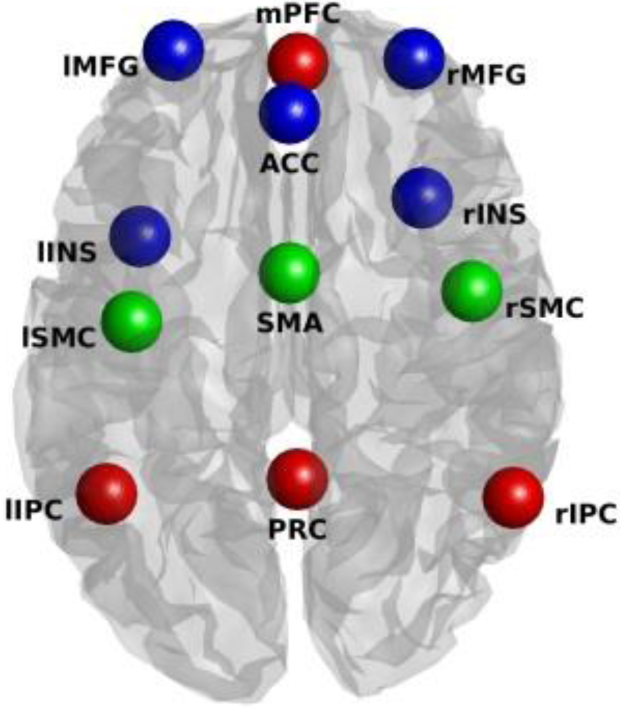
Location of regions included in the present study. Colors of spheres represent the network from which they were taken (blue represents salience network, red represents default mode network, and green represents somatomotor network). Anatomical labels: PRC = precuneus; l/rIPC = left/right intraparietal cortex; mPFC = medial prefrontal cortex; SMA = supplementary motor area; l/rSMR = left/right somatomotor region; l/rINS = left/right insula; ACC = anterior cingulate cortex; l/rMFG = left/right middle frontal gyrus.

#### 2.2.3 Dynamic Causal Modelling for resting state fMRI

Fully-connected DCMs without exogenous inputs were created and inverted for each session and subject separately (DCM12; DCM for resting state fMRI revision 6801). Four DCMs were estimated for each session: a DCM for each network separately and a DCM for the combined network. Default shrinkage priors – not informed by functional connectivity – were used for all networks. For each network (both separate and combined) sessions were excluded from further analyses if they did not meet all acceptance criteria (see also, Almgren et al., 2018) for both analysis types (i.e., with *and* without GSR). The acceptance criteria were: explained variance above 60%, at least one (extrinsic) connection strength greater than 1/8Hz, at least one effectively estimated parameter, maximum frame-wise displacement (FD) across all scans lower than 1.5mm, and a maximum alpha-threshold of 5% (i.e., type-I error rate; uncorrected for multiple comparisons) for which significant voxels were found in *all* regions of the specific network. For the longitudinal datasets; subjects were rejected if they had less than 8 sessions after diagnostic checks. This criterion excluded three subjects (i.e., S13, S18, and S19) for all networks, S14 was excluded for SMR, SAL and the combined network, and S4 was excluded for SMR. Additionally, 100, 77, 42, and 82 sessions were excluded (across subjects and datasets) for the SMR, SAL, DMN, and combined networks, respectively. For the HCP dataset, a total of 74, 65, 67, and 58 subjects were excluded because of failure to reach acceptance criteria for the SMR, SAL, DMN, and combined networks, respectively. Table 2 gives an overview of the sessions and subjects that were included for each dataset type.

**Table 2.** Number of included sessions and subjects for each network and dataset type. Abbreviations: HCP = Human Connectome Project, SMR = somatomotor network, SAL = salience network, DMN = default mode network; combined = combined network, subj = subjects.

| Dataset Type | Number sessions and/or subjects |  |  |  |  |  |  |  |  |  |
| --- | --- | --- | --- | --- | --- | --- | --- | --- | --- | --- |
|  | Total sample |  | SMR |  | SAL |  | DMN |  | Combined |  |
| Longitudinal | 653 sessions | (20 subj) | 499 sessions | (76%; 15 subj) | 533 sessions | (82%; 16 subj) | 578 sessions | (89%; 17 subj) | 528 sessions | (81%; 16 subj) |
| HCP | 361 subj |  | 287 subj | (80%) | 296 subj | (82%) | 294 subj | (81%) | 303 subj | (84%) |

#### 2.2.4 Parametric Empirical Bayes (PEB)

Connectivity at the subject or group level was modelled using a Parametric Empirical Bayesian (PEB; Friston et al., 2016) model, with a single regressor to model average connectivity across sessions or subjects. For the longitudinal datasets, two-level hierarchical PEB models were constructed. First, connectivity over sessions was estimated at the subject-level (i.e., average connectivity over sessions was computed for each subject separately). Then, subject-level PEB models were subsequently included in a group-level model; modelling average connectivity across subjects. Default PEB settings were used for estimation at the subject and group level (see, Almgren et al., 2018). Concerning the cross-sectional HCP dataset, connectivity was only estimated at the group level (i.e., across subjects), since only a single session was included for each subject. For both datasets, PEB models – equipped with a single between-session (or between-subject) precision component – were specified and estimated separately for connectivity and (spectral) noise. Spectral noise parameters represented three types of noise: global state noise (i.e., across-region neural fluctuations), global observation noise, and local (i.e., region-specific) observation noise.

#### 2.2.5 Global signal regression (GSR)

Dynamic causal modelling was performed with and without GSR. GSR was performed by adding a regressor representing the average signal intensity across the whole brain to the GLM model used to extract time-series (in addition to regressors representing motion, low-frequency fluctuations, white matter, and CSF signals). Time-series extracted from preprocessed images and were corrected for the effect of all regressors except the effect of low-frequency fluctuations. Analyses without global signal regression were performed in the same way, but excluding the regressor representing the average global signal.

#### 2.2.6 Assessing the quality of data features

In this work we wanted to compare the quality of *different data features* (i.e., time-series extracted with or without GSR), under the *same* model. For this, we adopted the Bayesian data comparison (BDC) approach described in Zeidman et al. (2019). This assesses the relative usefulness of data features (e.g., multiband factor of an fMRI acquisition scheme) in terms of making inferences about (group-level) parameters and models. The approach can be summarised as follows: usually, the approximate log model evidence (i.e., variational free energy), is used to assess the evidence for *different models* of the *same data*. The log evidence can always be decomposed into *accuracy* minus *complexity* (Penny, Stephan, Mechelli, & Friston, 2004). The complexity term represents the Kullback-Leibler divergence of the posterior density from the prior density. In other words, it scores the relative entropy or information gain afforded by the data. This is usually construed in terms of the number of degrees of freedom or parameters used to explain the data. However, it also reflects the reduction in uncertainty about model parameters after having seen the data. We therefore can use the decomposition of free energy into accuracy and complexity to ask whether different data features are more or less salient. In other words, we can quantify the reduction in uncertainty about model parameters afforded by one sort of data feature, relative to another – by simply looking at the differences in complexity. Generally speaking, we would consider data that were more informative (i.e., have a greater complexity) to be better than uninformative data. We therefore evaluated the complexity with and without GSR at the first level (i.e. after estimating DCMs at the session-level). To pool evidence across subjects, we assumed that the relative informative value of the data would be similar for all sessions and subjects. Therefor a fixed-effects approach was conducted to pool evidence across sessions (for the longitudinal datasets) and subjects (for the human connectome data). Practically, we computed the sum of the (log) complexities over sessions for both models for each subject separately. The difference between the present study and the empirical part of Zeidman et al. (2019) are the data features (i.e., data with or without GSR vs multiband factor) and the level at which usefulness of data was assessed (1^st^ level DCM vs group-level). Additionally, we also compared the certainty of posterior estimates for both analysis types.

## 3. Results

Detailed results for the longitudinal and cross-sectional datasets are outlined (separately) in subsections 3.1 and 3.2, respectively. Table 3 gives a summary of the main results.

**Table 3.** Summary of the results. Abbreviations: ^1^Root mean squared error, ^2^Hemispheric Asymmetry, ^3^Between-network, ^4^somatomotor network, ^5^default mode network, ^6^salience network.

| Influence of GSR on: | Longitudinal analysis | Cross-sectional analysis (HCP dataset) |
| --- | --- | --- |
| **Within-network connectivity**<br>(small RSNs) | **- Network-level:**<br>→ RMSD<sup>1</sup>: small ($\leq 0.1\text{Hz}$)<br>→ HA<sup>2</sup>: no difference in hemispheric dominance<br>**- Connection-level:**<br>→ negligible difference | **- Network-level:**<br>→ RMSD<sup>1</sup>: small ($\leq 0.1\text{Hz}$), except SMR (0.18Hz)<br>→ HA<sup>2</sup>: no difference in hemispheric dominance<br>**- Connection-level:**<br>→ some important differences, except for DMN |
| **Between-network (BN) connectivity** | **- Network-level:**<br>→ RMSD<sup>1</sup>: small (0.05Hz) | **- Network-level:**<br>→ RMSD<sup>1</sup>: acceptable ($\approx 0.1\text{Hz}$) |
|  | → Negligible influence on BN<sup>3</sup> connectivity | → Negligible influence on BN<sup>3</sup> connectivity |
|  | - **Connection-level:** | - **Connection-level:** |
|  | → Excitatory influences of SMR<sup>4</sup> and DMN<sup>5</sup> on SAL<sup>6</sup>, insignificant after GSR | → Excitatory influences became inhibitory after GSR (from SMR<sup>4</sup> and DMN<sup>5</sup> on SAL<sup>6</sup>) |
|  | → Additional inhibitory influence after GSR | → Additional inhibitory influences after GSR (from SAL<sup>6</sup> and SMR<sup>4</sup> on DMN<sup>5</sup>) |
| **Global observation noise parameters** | - No influence | - Significant influence, but not specific to observation noise parameters |
| **Optimal data features** | - **Small networks, separate estimates:** Data *without* GSR more informative, but less precise estimates<br>- **Combined network:** Data *without* GSR more informative, and more precise estimates | - **Small networks, separate estimates:** Data *without* GSR more informative, but less precise estimates<br>- **Combined network:** Data *without* GSR more informative, and more precise estimates |

### 3.1 Longitudinal Datasets

This section describes the results concerning the analysis using longitudinal datasets.

#### 3.1.1 The influence of GSR on within-network connectivity

Figure 2 shows the effect of global signal regression on effective connectivity within the three networks separately (i.e., SMR, SAL, and DMN). Estimated connectivity was remarkably similar for analyses with versus without GSR. *Across connections*, the average root mean squared difference (RMSD) between connectivity with and without GSR (self-connections converted to Hz) was 0.08, 0.05, and 0.04Hz for SMR, SAL, and DMN networks, respectively, which is smaller than the heuristic threshold of 0.1Hz usually used in DCM studies (see, e.g., Razi, Kahan, Rees, & Friston, 2015). On average, connectivity was slightly closer to the prior mean after GSR, however the effect was small (average difference in deviation from the prior mean was 0.06, 0.02, and 0.02Hz for SMR, SAL, and DMN, respectively).

**Figure 2.**
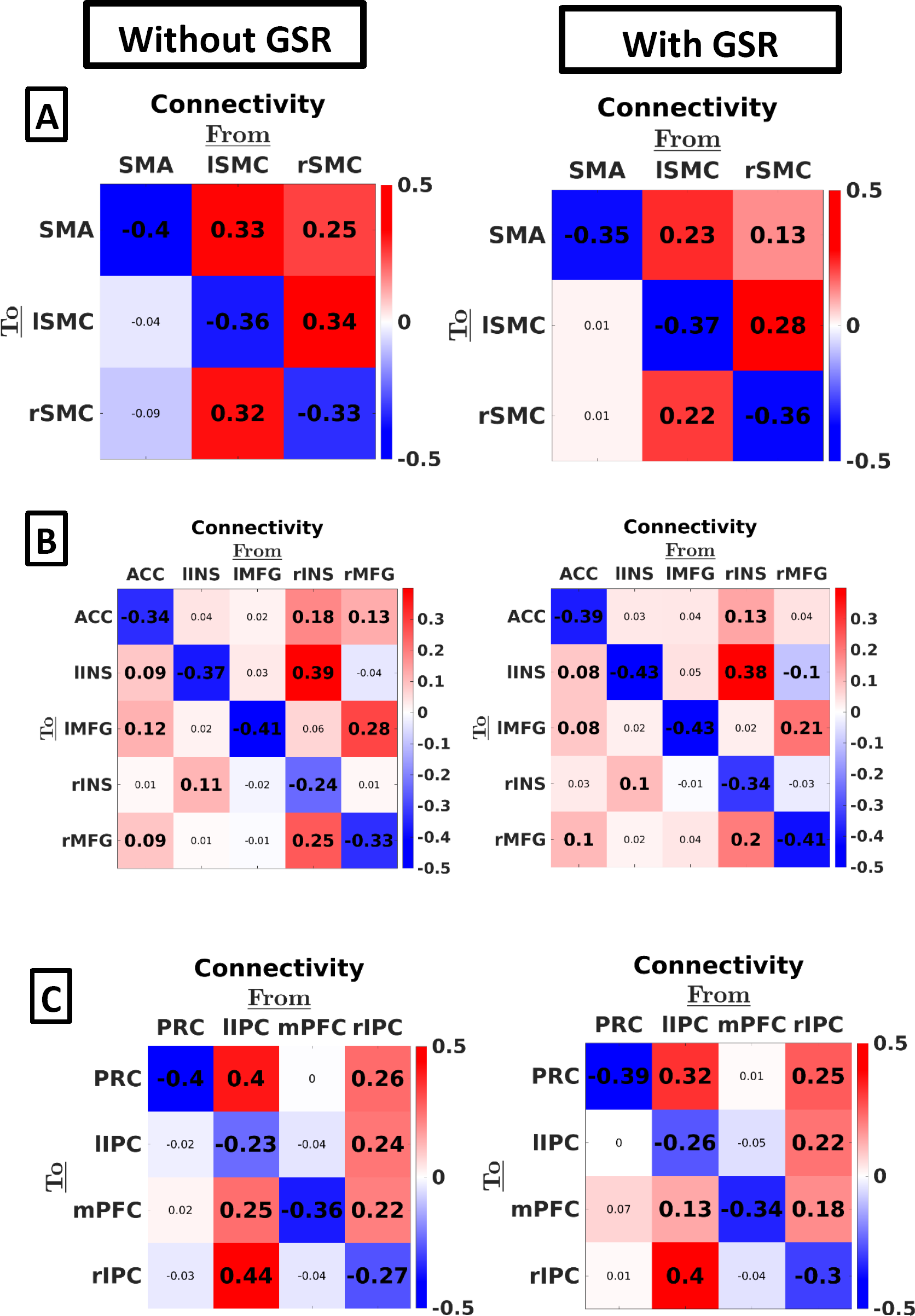
The effect of GSR on effective connectivity – using the longitudinal datasets – within three resting state networks: (A) SMR, (B) SAL, and (C) DMN. The color of the squares indicates the MAP connection strength for the respective connection (all in linear scale). Thresholding for inference was done using a posterior probability of 90%. Values in smaller fonts represent connections with a posterior probability smaller than 90%. Anatomical Abbreviations: SMA = supplementary motor area, l/r SMR = left/right somatomotor region, ACC = anterior cingulate gyrus, l/r INS: left/right insula, l/r MFG: left/right middle frontal gyrus, PRC = precuneus, l/r IPC = intraparietal cortex, mPFC = medial prefrontal cortex.

Global signal regression also had little effect on patterns of hemispheric asymmetry: the salience network showed higher outgoing influence from regions located in the right compared to the left hemisphere (without GSR: asymmetry = −0.14Hz, SD = 0.03Hz, posterior probability (PP) < 0.01 and with GSR: asymmetry = −0.08Hz, SD = 0.02Hz, PP < 0.01), while no hemispheric asymmetry concerning outgoing connectivity was found within the SMR in both cases (without GSR: asymmetry = 0.03Hz, SD = 0.06Hz, PP = 0.68; with GSR: asymmetry = 0.02Hz, SD = 0.05Hz, PP = 0.67). Results for asymmetry in the default mode network showed a higher influence from the left compared to the right hemisphere without and with GSR, although less evidence for asymmetry was observed for the latter (without GSR: asymmetry = 0.13Hz, SD = 0.05, PP = 0.99; with GSR: asymmetry = 0.07Hz, SD = 0.04Hz, PP = 0.93). In general, asymmetry was smaller for all networks after GSR.

*At the connection-level*, only three extrinsic connections (8% of a total of 38 extrinsic connections) showed a practically relevant change after GSR (i.e., difference > 0.1Hz; see also, Razi et al., 2015), namely the connection from lIPC to mPFC (part of DMN; see Figure 2: panel C), the connection from rSMC to SMA, and the connection from lSMC to rSMC (both part of the SMR network; see Figure 2: panel A). Additionally, an inhibitory connection emerged, and an excitatory connection disappeared after GSR (using a threshold of PP = 0.90).

#### 3.1.2 The influence of GSR on between-network connectivity

Figure 3 shows the effect of GSR on connectivity in the large combined network, which includes within- and between-network connectivity. Row A shows the connection-specific estimates, Row B shows the average connectivity between networks. Note the high similarity in within-network connectivity in the combined network compared to networks that were estimated separately (see, Figure 2).

**Figure 3.**
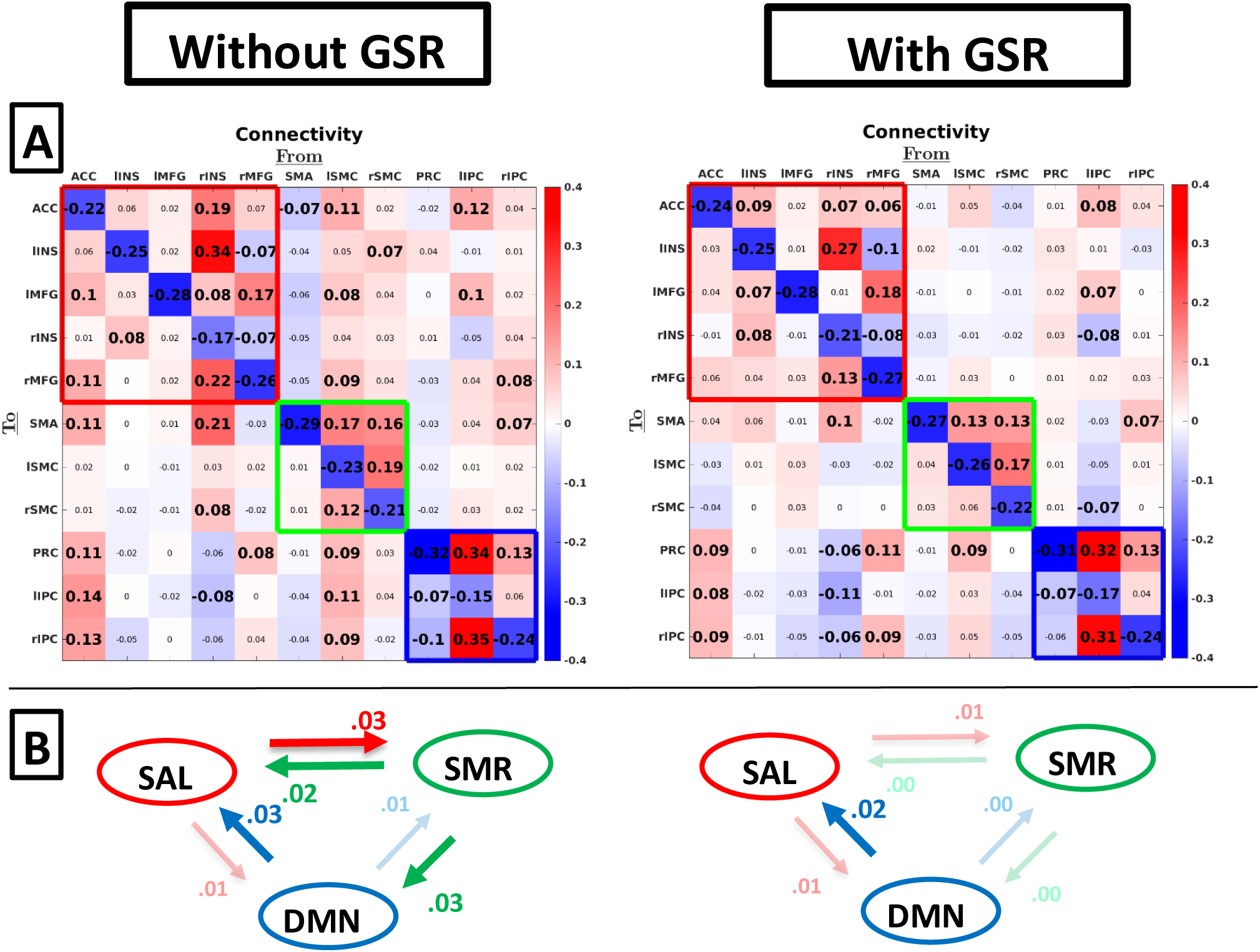
Group-level between-network connectivity without GSR (left column) and with GSR (right column). Row A shows estimates for individual connections. Red, green, and blue empty squares indicate connectivity within SAL, SMR, and DMN, respectively. Values in smaller fonts represent connections with a posterior probability smaller than 90%. Row B shows average connectivity between networks (across connections). Dimmed arrows and values depict connections with a posterior probability < 90%.

*Across between-network connections*, the effect of GSR was small: The root mean squared difference between connectivity with and without GSR was 0.05Hz across all between-network connections. Between-network connectivity was smaller in magnitude compared to within-network connectivity (difference in average between-network connectivity was 0.06Hz both with and without GSR, excluding self-connections). The mean influence networks had on each other (panel B) was negligible (< 0.05Hz) both with and without GSR.

For *individual connections*, we found small excitatory reciprocal connectivity between all three networks without GSR, with weak inhibitory connectivity from rINS to lIPC and from SMA to ACC. After GSR reciprocal connectivity between SAL and SMR networks disappeared, and weak inhibitory influence was found from rINS (salience network) on all regions of the DMN. The ACC had an excitatory influence on the DMN both with and without GSR.

#### 3.1.3 The influence of GSR on noise parameters

To have sufficient spatially distributed information to allow robust estimates of (global) spectral noise components – and their average across sessions or subjects – we only analysed the combined network here. The noise here is modelled by using power law distribution having two parameters – amplitude and an exponent – to characterize their cross spectral density. Figure 4 shows the effect of GSR on parameters representing (spectral) noise, which include endogenous fluctuations that drive the system (i.e., state-noise) and global measurement (or observation) noise. The first parameter of each noise-component represents the amplitude of fluctuations, while the second parameter represents the shape (exponent) of the noise spectrum. Clearly, GSR did not have an effect on any of the (global) noise components.

**Figure 4.**
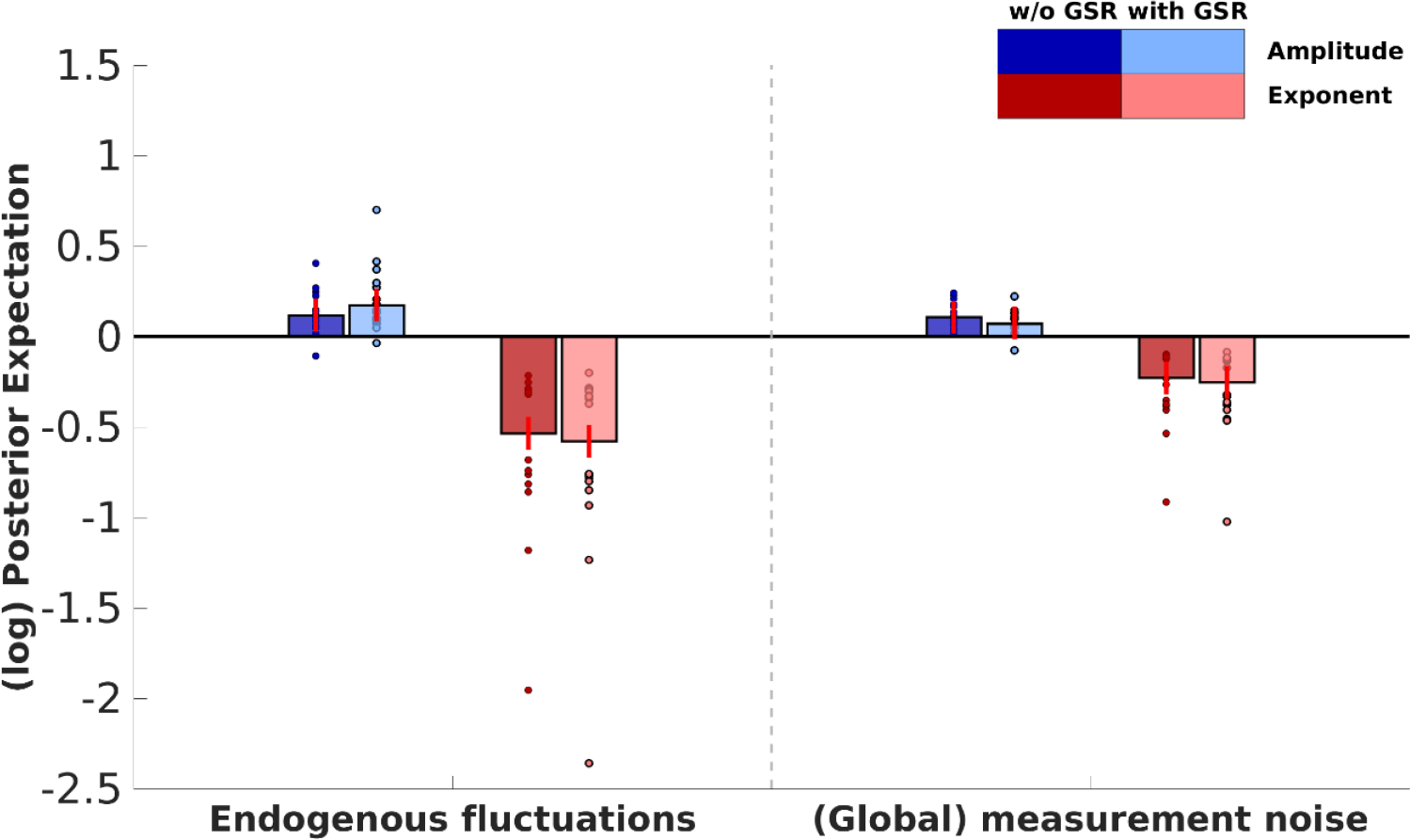
The effect of GSR on (spectral) noise parameters within the combined network. Bar heights depict the group-level maximum a posteriori (MAP) estimates for the respective parameter, small circles depict subject-specific estimates (not empirically optimized). The color of the bars depict the parameter type (i.e., amplitude and frequency). Lightness of the bars and markers depicts connectivity without or with GSR (darker and lighter, respectively). Red lines depict 90% credibility intervals for the posterior estimates (PEB.Cp). Parameters are shown in log-scale and are relative to the prior mean.

#### 3.1.4 Identifying data features

To identify the most informative data features, we decomposed free energy into accuracy and complexity, where accuracy is the expected likelihood of the data under posterior beliefs and complexity is the Kullback-Leibler divergence between the posterior and prior densities. Complexity can thus be considered as a measure of the informative value of the data (Zeidman et al., 2019). Therefore, we compared the estimated complexity with and without GSR at the session-level, which is directly dependent on the data. We used a fixed-effect approach to pool complexity over sessions for each subject separately (see, e.g., Stephan, Penny, Daunizeau, Moran, & Friston, 2009). DCM estimation *without* GSR was found to have the highest complexity in 93.3% of subjects for the SMR network, in 94.0% of subjects for the SAL network, 88.2% of participants for the DMN, and for 100% of subjects in the combined network, with strong evidence in all but one subject (Bayes factor greater than 20; Kass & Steffey, 1989). However, the certainty of posterior estimates (computed as the negative entropy of the posterior distribution) was greater for the data with GSR in 60.0%, 82.4%, 87.5% of subject for SMR, DMN, SAL, respectively. For the combined network, the opposite pattern was observed: 81.2% of subjects showed more precise estimates without GSR.

### 3.2 Cross-sectional Dataset

This section outlines the results concerning the HCP dataset.

#### 3.2.1 The influence of GSR on within-network connectivity

Figure 5 shows estimated effective connectivity in the three resting-state networks, both with and without GSR. *Across all connections*, the average root mean squared difference (RMSD) between connectivity with and without GSR was 0.18, 0.10, and 0.07Hz for SMR, SAL, and DMN networks, respectively. The average decrease in deviation from the prior mean with GSR (compared to without GSR) was 0.08, 0.04, 0.05Hz for SMR, SAL, and DMN, respectively. Global signal regression had little effect on patterns of hemispheric asymmetry: the salience network showed higher outgoing connectivity from regions located in the left compared to the right hemisphere (without GSR: asymmetry = 0.10Hz, SD = 0.02Hz, PP > 0.99; with GSR: asymmetry = 0.06Hz, SD = 0.01Hz, PP > 0.99), right hemispheric dominance was found within the SMR for both cases (without GSR: asymmetry = – 0.47Hz, SD = 0.04, PP < 0.01; with GSR: asymmetry = −0.16Hz, SD = 0.03Hz, PP < 0.01), and left dominance was found for the DMN with and without GSR (without GSR: asymmetry = 0.15Hz, SD = 0.03Hz, PP > 0.99; with GSR: asymmetry = 0.11Hz, SD = 0.02Hz, PP > 0.99). As with the analysis of longitudinal designs, asymmetry decreased for all networks after GSR.

**Figure 5.**
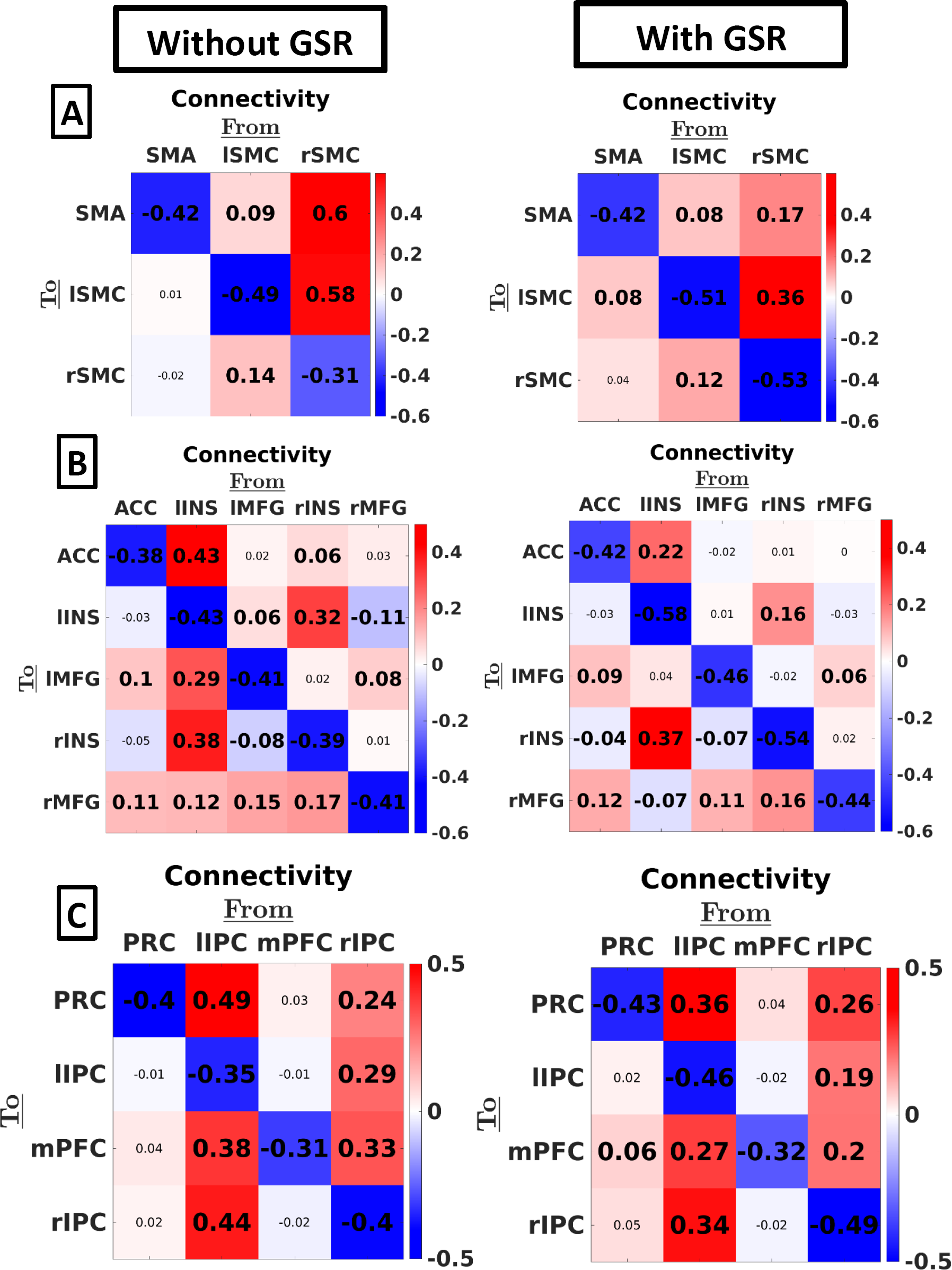
The effect of GSR on effective connectivity – using the HCP data – within three resting state networks: (A) SMR, (B) SAL, and (C) DMN. The color of the color of the squares indicates the MAP connection strength for the respective connection (all in linear scale). Thresholding for Inference was done using a posterior probability of 90%. Values in smaller fonts represent connections with a posterior probability smaller than 90%. Anatomical Abbreviations: SMA = supplementary motor area, l/r SMR = left/right somatomotor region, ACC = anterior cingulate gyrus, l/r INS: left/right insula, l/r MFG: left/right middle frontal gyrus, PRC = precuneus, l/r IPC = intraparietal cortex, mPFC = medial prefrontal cortex.

*At the connection-level*, nine extrinsic connections (24% of extrinsic connections) showed a practically significant change after GSR (i.e., difference > 0.1Hz): two connections in SMR, four connections in SAL, and three connections in DMN. In all except two connections these were outgoing connections from the (network-specific) dominant region. Seven extrinsic connections (18%) emerged or disappeared (threshold PP = 0.90) after GSR (although they were all small in magnitude). Additionally, one extrinsic connection changed sign (from excitatory to inhibitory or *vice versa*) after GSR.

#### 3.2.2 The influence of GSR on between-network connectivity

Figure 6 shows the between- and within network connectivity for the combined network. Row A shows the separate connectivity estimates, Row B shows the average connectivity between networks.

**Figure 6.**
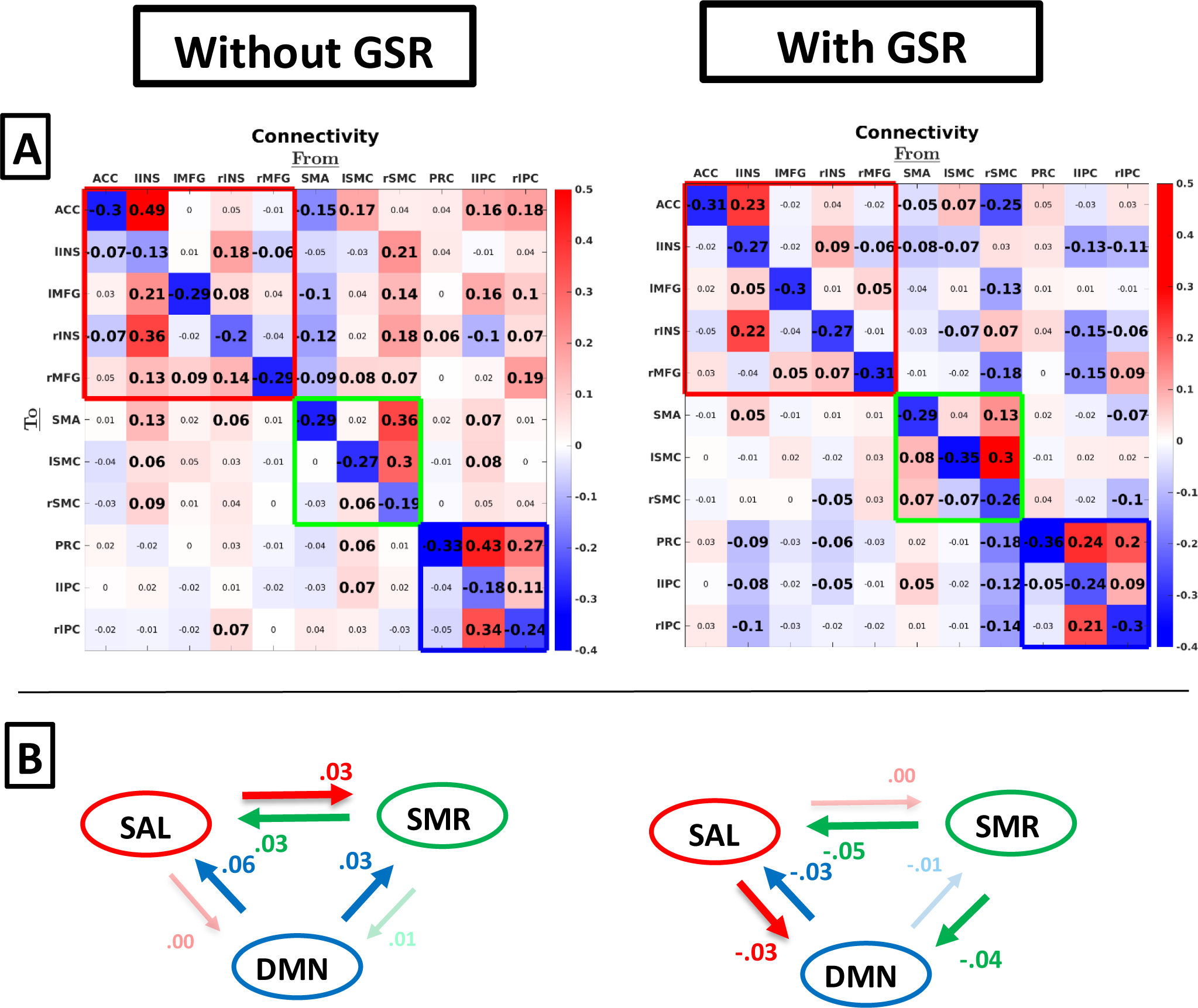
Group-level between-network connectivity without GSR (left column) and with GSR (right column). Row A shows estimates for each connection separately. Red, green, and blue empty squares indicate connectivity within SAL, SMR, and DMN, respectively. Values in smaller fonts represent connections with a posterior probability smaller than 90%. Row B shows average connectivity between networks. Dimmed arrows and values depict connections with a posterior probability < 90%.

*Across connections*, the root mean squared difference between connectivity with and without GSR was 0.10Hz across all between-network connections, with larger within-network connectivity compared to between-network connectivity (difference in mean magnitude of connection was 0.08Hz without GSR and 0.03Hz with GSR, excluding self-connections). Note again the striking similarity in connectivity compared to the separately estimated networks (see, Figure 5). The average influence networks had on each other (Figure 6; Row B) was negligible irrespective of processing method, although some small excitatory influences became slightly inhibitory after GSR.

*At the level of individual connections*, the influence of GSR was larger compared to the average between-network connectivity: without GSR we mainly observe excitatory influence from SMR and DMN on SAL, which becomes mainly inhibitory after GSR (although some inhibitory connectivity from SMA on most regions of SAL was found even without GSR). Additionally, after GSR we observe inhibitory connectivity from SAL on DMN, and from rSMC on all DMN regions, which was not found without GSR.

#### 3.2.3 The influence of GSR on noise parameters

Figure 7 shows the effect of GSR on parameters representing (spectral) noise, which include endogenous fluctuations that drive the system (i.e., state-noise) and global measurement (or observation) noise. Clearly, GSR had an effect on both estimated endogenous fluctuations and global measurement (observation) noise, showing that the effect of GSR is not specific to the parameter representing global measurement noise. Note that the exponent components (i.e. parameters that model the shape of the spectrum) of group-effects yielded in some cases unintuitive results (i.e., group estimates at lower end of subject-specific estimates), which most likely arise from complex covariance structures between these components (see, Kasess et al., 2010 for similar unintuitive results using Bayesian parameter averaging in DCM).

**Figure 7.**
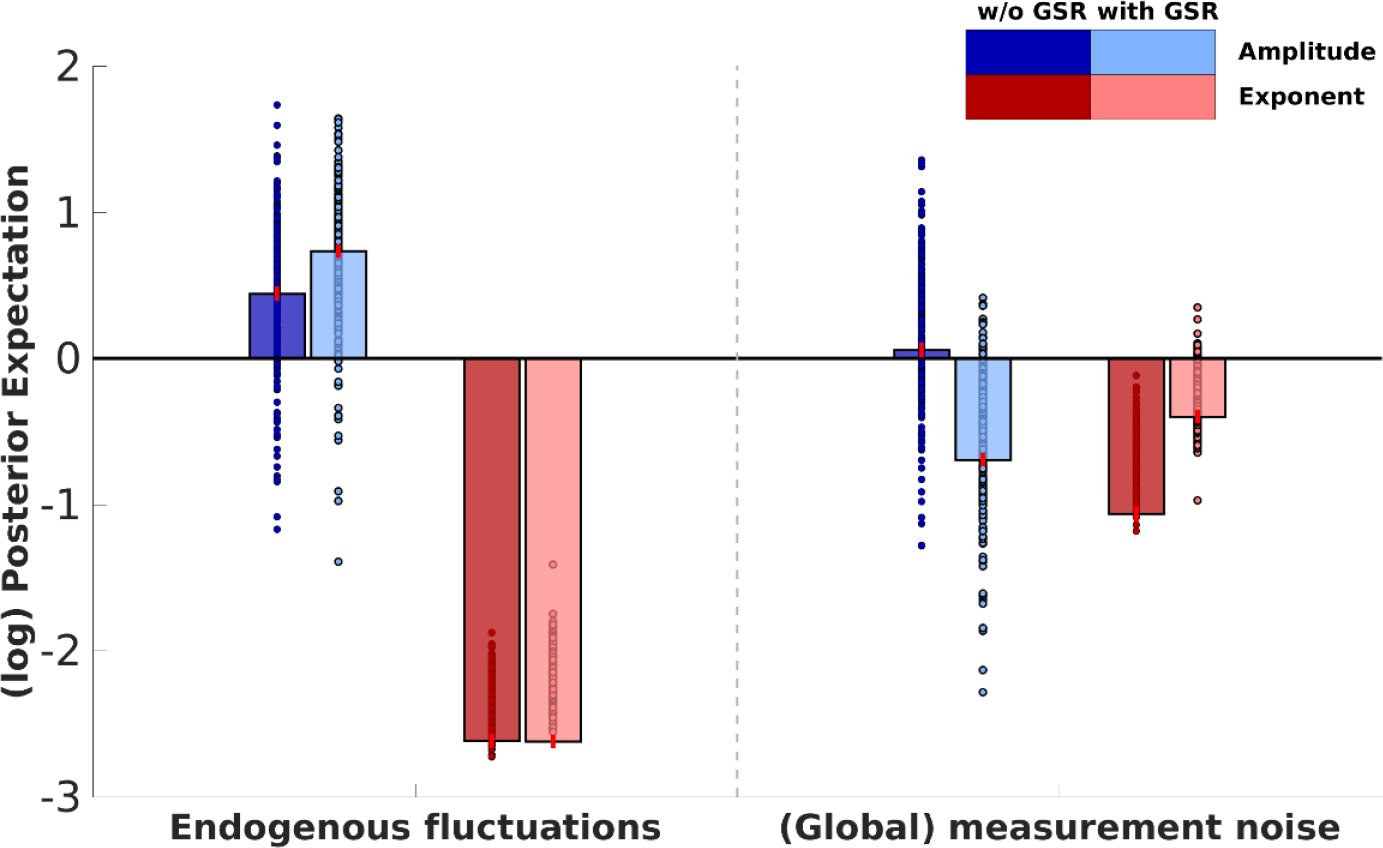
The effect of GSR on (spectral) noise parameters within the combined network. Bar heights depict the group-level maximum *a posteriori* estimates for the respective parameter, small circles depict subject-specific estimates. The color of the bars depict the parameter type (i.e., amplitude and exponent). Lightness of the bars and markers depicts connectivity without or with GSR (darker and lighter, respectively). Red lines depict 90% credible intervals for the posterior estimates (PEB.Cp). Parameters are shown in log-scale and are relative to the prior mean.

#### 3.2.4 Identifying data features

A fixed-effect approach was used to pool complexity over subjects (see, e.g., Stephan, Penny, Daunizeau, Moran, & Friston, 2009). For all networks, DCM estimation without GSR was found to have greater complexity – or information gain – compared to estimation with GSR, with strong evidence in all cases (Bayes factor greater than 20; Kass & Steffey, 1989). However, again the certainty of the estimates (i.e., negative entropy) was greater for the data with GSR for SMR, DMN, SAL, respectively, but the opposite pattern was found for the combined network.

## 4. Discussion

In this study we investigated the effect of GSR on effective connectivity and parameters representing global observation noise, estimated with spectral DCM. Additionally, we investigated which data (i.e., data extraction with or without GSR) were most informative (i.e., have the largest complexity) for estimation of DCM parameters. We focused on both within- and between-network connectivity in two different designs (longitudinal and cross-sectional). The use of different designs allowed us to investigate the generalizability of our conclusions, and to discover divergent effects of GSR in different designs.

In general, we found negligible to small effects of GSR on connectivity within small (separately estimated) RSNs. At the network-level the effect was very small, both in terms of root-mean-squared deviation as well as hemispheric asymmetry. Additionally, the majority of individual connections did not change sign. These results show that GSR in small RSNs should not constitute a major concern in future research using resting state DCM. Our results agree with studies of functional connectivity, showing that connections with *functionally related* areas mostly remain significantly positive after GSR (e.g., Chang & Glover, 2009; Weissenbacher et al., 2009). However, a few connections became inhibitory after GSR, which was more observable in the HCP dataset. Although these effects were fairly small, they might be important to consider in future research.

Concerning between-network connectivity, we found two important effects: the effect of GSR on the between-network connectivity *across* connections (i.e., net influence between networks and root-mean-squared difference) was negligible to small, while the effect of GSR on *individual* connections was non-negligible. At the level of individual connections many connections between DMN and SAL were excitatory when GSR was not applied, most prominently for the HCP dataset (unidirectional influence from DMN in case of HCP), while most connections were inhibitory (and slightly more bidirectional) *after GSR*. Furthermore, we found both excitatory and inhibitory influence from SMR on SAL, which became mainly inhibitory or non-existent after GSR. The general decrease in excitatory (and increase in inhibitory) between-network connectivity after GSR has also been found in studies focusing on functional connectivity (see, Murphy & Fox, 2017). In conclusion, the robustness of network-level effects, in contrast to connection-level effects, with respect to GSR shows that DCM connections are best interpreted jointly (which has been argued in other studies, e.g., Almgren et al., 2018).

The between-network results are somewhat in contrast to the DCM study of Zhou et al. (2018), which found inhibitory influence from the salience network on the core DMN, but not *vice versa, without* GSR (which we found in the opposite direction after GSR in the HCP dataset). There are three important differences in results between Zhou et al. (2018) and the result we obtained in the present study that should be explained. First, Zhou et al. found inhibitory between-network connectivity in the absence of GSR. This is probably explained by their use of a wide set of motion-related regressors (24 regressors coding both instant as preceding motion, as well as their squares). Since GSR is closely linked to motion (Liu et al., 2017), these motion parameters might have a similar effect as GSR, hence excluding spurious positive connectivity. Second, for the HCP data we found mainly inhibitory connectivity (after GSR) from DMN to SAL, while Zhou et al. (2018) found inhibitory connectivity in the opposite direction. This might be explained by a difference in subject characteristics: Subjects in our study were mainly young adults (mean age was 30.1 and 28.7 years respectively, for longitudinal and cross-sectional design), while the age-group included in Zhou et al. (2018) was on average younger (mean age = 17.4 years). However, note that very small inhibitory connectivity from specific regions of SAL on DMN without GSR, but this did not reach our threshold (posterior probability < 90%). Third, Zhou et al. (2018) found important between-network connectivity when averaged across connections, while we only found effects at the connection-level (see, e.g., Figure 6). This might be explained by their use of functional connectivity to inform priors: between-network effects might be more coherent when connectivity parameters are informed by FC, hence increasing detection of between-network effects. In our study we did not use FC-informed priors for two reasons: (1) Bayesian model comparison showed strong evidence in favour of the use of standard (i.e., shrinkage) priors, (2) the use of group-specific (across-subject) FC-priors at higher levels in our hierarchical design might have obscured results. Additionally, the difference in sample size and specific ROI specification method might also have had an effect on the divergent results.

On average, connectivity was closer to the prior mean after GSR. Moreover, hemispheric asymmetry (difference between left and right outgoing connectivity) also decreased after GSR. For extrinsic connectivity, this is in line with multiple studies showing that the connectivity decreases after GSR (e.g., Chang & Glover, 2009; Fox et al., 2009; Weissenbacher et al., 2009). The deviation from the prior mean has also influence on the informative value of the data, which is discussed further in this discussion. Notably, hemispheric asymmetry also decreased after GSR. Hemispheric asymmetry was here defined as the (network-specific) outgoing connectivity from regions in one hemisphere relative to the outgoing connectivity from regions in the other hemisphere. Hence, a change in hemispheric asymmetry would mean that the magnitude of the decrease in connectivity is different for regions in both hemispheres. Possibly the extent to which certain regions contribute to the global signal differed between hemispheres. This explanation is in line with McAvoy et al. (2016) who found an asymmetric contribution of regions to the global signal. However, a direct comparison with the functional connectivity literature concerning the size of the effect of GSR is difficult given the differences in models, difference in statistical framework (Bayesian vs frequentist), and specific networks under investigation. It is also important to note that our conceptualization of hemispheric asymmetry differs from that in functional connectivity studies, since DCM estimates directed influences.

Remarkably, the *within-network* connectivity was very similar when networks were estimated separately compared to networks as a whole (compare e.g., Figure 5 and 6). This was also found by Ushakov et al. (2016), who showed that connectivity within the core DMN was very similar when either left or right hippocampus was included in the network. This shows that group-level resting-state DCM results are quite robust against addition of extra networks and regions.

Contrary to our expectations, we either found no effect (longitudinal designs) or an unspecific effect (cross sectional design) on measurement noise parameters. Several explanations can be put forward to account for these counter-intuitive results, and the explanations might be different between both designs. Concerning the absence of effect in the longitudinal designs, it might be that the effect of GSR was captured by the between-subject components of the hierarchical Bayesian model. This is possible, since the datasets differed in terms of scanner type, pulse sequence parameters, and subject characteristics, which might cause divergent effects on noise parameters. Therefore, it is possible that the effect of GSR was cancelled out at the group-level, while captured by parameters representing between-subject variance. Indeed, we found an important effect (i.e., increase vs decrease in parameters) of GSR on noise parameters for some individual subjects, but not for others. Concerning the cross-sectional HCP dataset, the effect was captured by multiple noise parameters. This unspecific effect is possible, since resting state fMRI has no explicit exogenous input (e.g., an experimental design) that could inform more precise estimation. Noise parameters that are shared across regions (e.g., global state and measurement noise) might therefore be all sensitive to the global signal.

Additionally, we asked whether data was more informative for 1^st^ level parameter estimation (i.e., had the highest complexity) with or without GSR. Connectivity estimation using data without GSR was found to yield the greatest complexity for all datasets and networks. The complexity term is the Kullback-Leibler divergence between the prior and posterior density, which can be interpreted as the increase in information after updating the prior with the data. Complexity essentially depends on (1) divergence of the mean of the posterior from the mean of the prior density, (2) precision of the posterior density compared to the precision of the prior, and (3) change in covariance patterns between posterior and prior densities of parameters. The observations of higher complexity without GSR in most cases probably is for a great part attributable to a greater divergence from the posterior mean. The few (subject-specific) observations of higher complexity with GSR are at odds with this explanation, since after GSR a smaller deviation from the prior mean was observed. However, the precision of the parameter estimates also has an influence on the KL-divergence. Therefore, we compared the precision of estimations (computed as the negative entropy) for data with versus without GSR. Here we found that for *small networks*, data with GSR was found to yield greatest precision. However, for the combined network, data *without* GSR was found to yield most precise estimates. This might be associated with the (potentially spurious) inhibitory connectivity that was found for estimation with GSR in the combined network.

In general, we can thus conclude that data without GSR are more informative for estimation of effective connectivity. One should note that this complexity was computed at the session-level, and that this level is directly dependent on the data. For the other levels (i.e., subject- and group-level), the complexity term includes the Kullback-Leibler divergence of second-level parameters (e.g., between-session variance; group-level effective connectivity) from their prior density, which depends on the estimated parameters at lower levels, and not directly on the data. Possibly, between-session (between-subject) differences and subject (group) level estimates are less informed by the estimates based on data without GSR.

In this work we assessed the (practical) effects of GSR on parameters estimated with DCM for resting state fMRI. However, multiple alternative methods exist to eliminate (global) noise from fMRI data, including temporal ICA (Glasser et al., 2018), dynamic global signal regression based on blood arrival time (dGSR; Erdoğan, Tong, Hocke, Lindsey, & deB Frederick, 2016), and corrections for respiratory and cardiac signals (Chang, Cunningham, & Glover, 2009). Although these alternatives are less extensively studied in fMRI research compared to ‘canonical’ GSR, they have certain features that *might* make them superior to GSR. These methods are outside of the scope of the present study, which was to assess the effect of the (widely studied) ‘canonical’ GSR on DCM parameters. However, we also checked the consistency of our results compared to the effects of global signal *normalization* (dividing timeseries by the global signal instead of regressing out), and we found quite similar results.

In conclusion, GSR is a minor concern in DCM studies. However, individual between-network connections (as opposed to average between-network connectivity) should be interpreted with some caution. Additionally, we suggest the use of the complexity term – in combination with the certainty of estimation – to assess the relative informative value of the data with versus without GSR.

## Software availability

Code to reproduce analyses, results and figures is available at: https://github.com/halmgren/Pipeline_effect_GSR_effective_connectivity_rsfMRI.

## Acknowledgements

This work has been funded by the Special Research Fund of Ghent University (awarded to HA; grant No. BOF16/DOC/282; https://www.ugent.be/), a grant for long stay abroad from the Fund for Scientific Research-Flanders (FWO-V; awarded to HA; http://www.fwo.be), the Fund for Scientific Research-Flanders (FWO-V; grant No. FWO14/ASP/255 awarded to FvdS; http://www.fwo.be), the Australian Research Council DECRA Fellowship (awarded to AR; Ref: DE170100128), and funds from the Wellcome Trust (KF, AR).

## Notes

https://github.com/halmgren/Pipeline_effect_GSR_effective_connectivity_rsfMRI

